# Acidification-dependent suppression of *C. difficile* by pathogenic and commensal enterococci

**DOI:** 10.1101/2023.05.16.541032

**Authors:** Holly R. Neubauer, Ibukun M. Ogunyemi, Alicia K. Wood, Angus Johnson, Avi Z. Stern, Zainab Sikander, Lesly-Hannah Gutierrez, Addelis A. Agosto, Peter T. McKenney

## Abstract

*Clostridioides difficile* and Vancomycin-resistant *Enterococcus faecium* (VRE) are commonly co-isolated from hospitalized patients. We sought to develop a co-culture biofilm model to characterize interactions between these two opportunistic pathogens. Upon growth in biofilm-promoting media containing added glucose, fructose or trehalose, VRE produces sufficient acid to lower the pH and inhibit growth of *C. difficile*. We found this effect depended on the carbon source, and that acidification by VRE was necessary and sufficient to suppress *C. difficile* growth in liquid medium and in cecal content extracts from germ free mice. VRE frequently dominates the intestine of patients administered antibiotics which can predispose to the development of *C. difficile* infection. We reasoned that it may be possible to suppress *C. difficile* growth during co-infection with VRE by supplementing the mouse diet with a fermentable sugar. A VRE-dominated gut microbiota may convert the sugar to acid, lower the pH and reestablish colonization resistance to *C. difficile*. Supplementation of the diet of VRE colonized mice with high levels of fructose neither resulted in a lower pH, nor did it prevent colonization by *C. difficile*. Taken together, these data suggest that VRE can suppress growth of *C. difficile* by organic acid production in a carbon source-dependent manner in vitro, however, the mammalian intestine may require sophisticated approaches to lower pH therapeutically.

## Introduction

High population diversity in the gut microbiota is generally correlated with health across gastro-intestinal diseases. Treatment with antibiotics can disrupt the gut microbiota to the point that a single species can expand and dominate the gut, making up a high percentage of the reads in a 16S microbiome sequencing data set. Treatment with broad-spectrum antibiotics can lead to expansion of VRE (Donskey et al., 2000) and domination of the gut microbiota by the genus *Enterococcus*. This was observed in humans (Taur et al., 2012) and mice (Ubeda et al., 2010). This domination occurs in hematopoietic stem cell transplant patients where a low diversity microbiota is correlated with mortality (Liao et al., 2021; Taur et al., 2012). Vancomycin-resistant *Enterococcus faecium* (VRE) is a rising clinical concern that causes difficult-to-treat systemic infections. But it is often forgotten that *E. faecium* is also a common member of the human gut microbiota, a lactic acid bacterium, and a commonly used probiotic in humans and agricultural animal production (Dubin & Pamer, 2014; Krawczyk et al., 2021; Yahan Wei et al., 2024).

The majority of studies suggest that patients with a low-diversity gut microbiota following antibiotic treatment are also at elevated risk for *Clostridioides difficile* infection (CDI). Colonization of the gut with VRE and the genus *Enterococcus* has been correlated with increased risk of CDI in hospitalized human patients (Fujitani et al., 2011; Lee et al., 2017; Smith et al., 2022). *Enterococcus* species in the gut microbiota were predicted to enhance CDI risk in mathematical modeling that combined data from human and mouse infections (Buffie et al., 2015). In mouse models, genus *Enterococcus* was also correlated with increased persistence of *C. difficile* in the gut (Tomkovich et al., 2020) and increased toxin production and virulence under high dietary zinc conditions (Zackular et al., 2016). One other study described correlation of genus *Enterococcus* abundance with attenuation of *C. difficile* virulence (De Wolfe et al., 2019). It is possible that strain diversity may contribute to differing results as some commensal enterococci suppress *C. difficile* growth *in vitro* (Rolfe et al., 1981).

Two direct tests of the effects of *Enterococcus* colonization on the CDI mouse model reported higher toxin titer and pathology (Keith et al., 2020; Smith et al., 2022). The mechanism of enhancement of CDI severity in mice was linked to metabolic cross-talk between *E. faecalis* OG1RF and *C. difficile* in which arginine supplementation was sufficient to reduce toxin production and pathology, without significantly affecting colonization by either organism (Smith et al., 2022). Furthermore, *E. faecalis* benefits from the release of host heme caused by *C. difficile* toxin-mediated damage in mouse models of CDI (Smith et al., 2024).

Both *C. difficile* and enterococci form biofilms (Ch’ng et al., 2019; Tremblay & Dupuy, 2022). In both species, biofilms contribute to antibiotic tolerance (Dapa & Unnikrishnan, 2013; Holmberg & Rasmussen, 2016) and may be a reservoir for infection recurrence in CDI (Frost et al., 2021; Normington et al., 2021). This study began as an attempt to establish an in vitro co-culture biofilm model of VRE and *C. difficile*. We quickly noticed that the byproducts of primary metabolism produced by VRE negatively affected *C. difficile* growth. Here we have established that VRE and commensal enterococci are capable of acidifying unbuffered bacterial growth media below a pH that is growth inhibitory to *C. difficile* and other commensal and opportunistic clostridia. Under these conditions, we show that acidification of a carbon source is necessary and sufficient for inhibition of *C. difficile* and clostridia growth. These data suggest that in conditions where lactic acid bacteria such as VRE dominate the local environment, alteration of acid production and pH levels in a carbon source-dependent manner affect the viability of *C. difficile* in co-culture *in vitro*. Finally, we modeled the effects of carbon source control in VRE dominated mice by supplementing the diet with high levels of fructose followed by *C. difficile* infection. Fructose supplementation did not affect *C. difficile* CFU levels in VRE co-infected mice. In our mouse model of co-infection, fructose supplementation alone was not sufficient to alter *C. difficile* colonization, suggesting that more complex interventions may be necessary to reproduce the in vitro phenomenon.

## Results

### Glucose metabolism inhibits *C. difficile* growth in co-culture with VRE

We began this work as an attempt to establish a dual-species in vitro liquid biofilm model for VRE and *C. difficile*. Typically for Gram-positive bacteria, glucose is added to the culture media to a final concentration of 0.2-1%, which promotes attachment and production of exopolysaccharide matrix (Dapa & Unnikrishnan, 2013; Donelli et al., 2012; Kristich et al., 2004; Pillai et al., 2004; Toledo-Arana et al., 2001). When we performed a pilot experiment of liquid co-culture growth in Supplemented Brain Heart Infusion (BHI) broth with 0.4% added glucose (total glucose = 0.6%) we found a significant decrease in *C. difficile* growth at 8 and 24 hours post-inoculation when in co-culture with VRE (Figure 1A). We observed a similar elimination of *C. difficile* when co-cultured with VRE in Sporulation Media (SM) which lacks an added carbon source, when it was supplemented with 0.4% glucose or greater in co-culture (Figure 1C).

**Figure 1.**
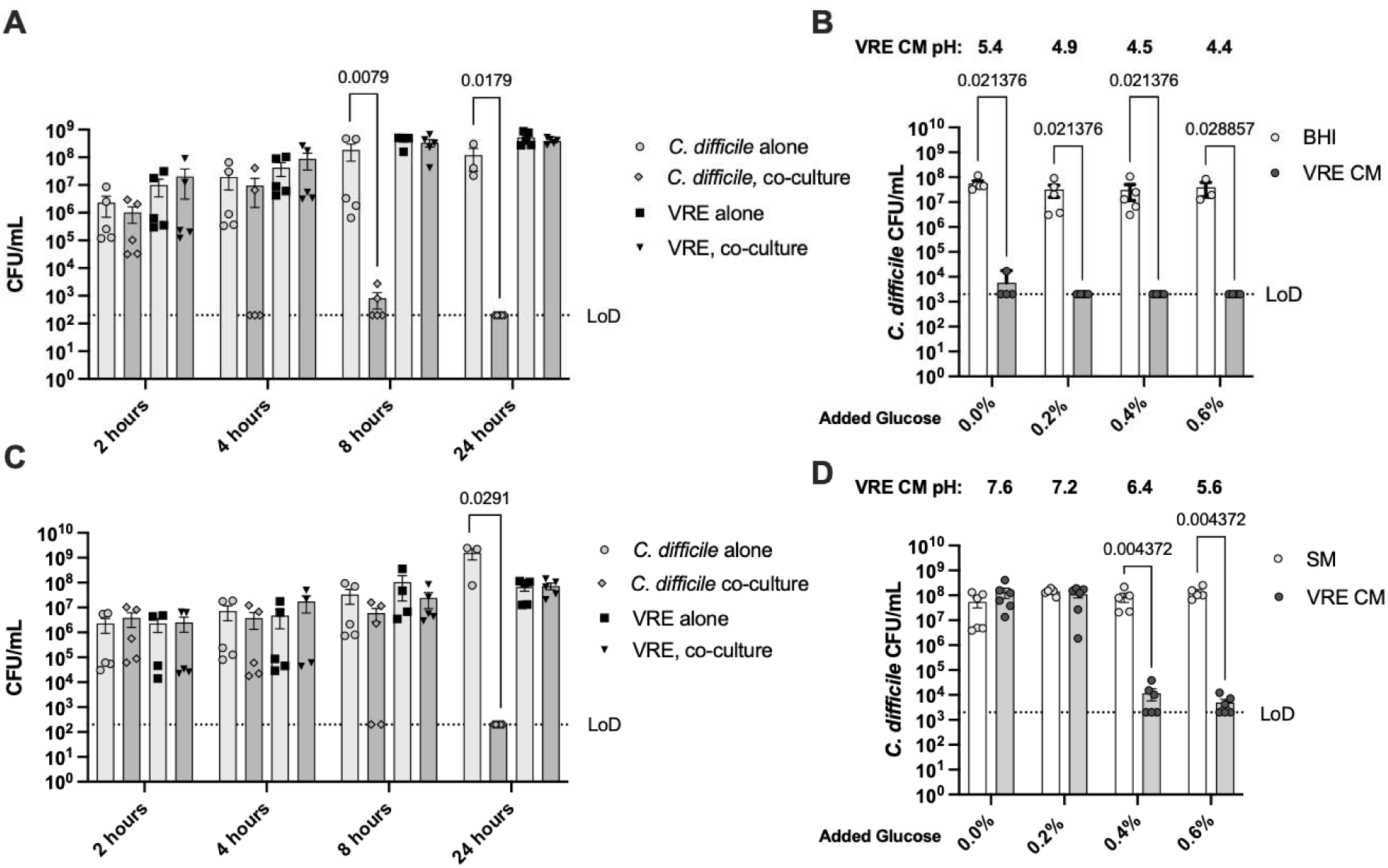
*C. difficile is* inhibited by VRE during co-culture in excess glucose. A) Time course CFUs of *C. difficile* and VRE alone and in co-culture in BHI + 0.4% glucose (0.6% total glucose). Data are combined from 2 independent experiments, n = 3-5 biological replicates per time point, Mann-Whitney test of alone vs. co-culture. B) VRE was grown for 48 hours in BHI + 0, 0.2, 0.4 or 0.6% additional glucose, then filter sterilized to create conditioned media (VRE CM) which was then inoculated with *C. difficile*. The average pH of the VRE CM is shown above the graph. Data are combined from 2 independent experiments, n = 3-5 biological replicates per time point, Mann-Whitney test. C) Time course CFUs of *C. difficile* and VRE alone and in co-culture in SM + 0.6% glucose. Data are combined from 2 independent experiments, n = 3-5 biological replicates per time point, Mann-Whitney test of alone vs. co-culture. D) VRE was grown for 48 hours in SM + 0, 0.2, 0.4 or 0.6% additional glucose. CM was filter sterilized to create VRE CM, the average pH of the VRE CM is shown above the graph. It was then inoculated with mid-log *C. difficile* and grown for 48 hours. Data are combined from 2 independent experiments, n = 5-6 biological replicates per time point, Mann-Whitney test.

Next to determine if the inhibitory factor produced by VRE is soluble and filterable, we generated conditioned media by growing VRE to exhaustion (OD600 ∼ 1.8) and filtered it through a 0.22 μm filter. To determine if the inhibition was dose-dependent, we generated VRE conditioned media with escalating levels of added glucose. The conditioned media was then inoculated with *C. difficile* and grown for 48-hours. We found that increasing the concentration of glucose results in inhibition of *C. difficile* growth and that the inhibitory factor is soluble and filterable in both BHI and SM (Figure 1B & 1D). Enterococci are closely related to lactic acid bacteria and are known to produce large amounts of acid when cultured in glucose (Ramsey, Hartke, and Huycke 2014). We tested the pH of the VRE conditioned media and found that pH is inversely correlated with glucose concentration. In BHI conditioned media the starting pH was < 5.3 for all tested glucose concentrations, which did not support *C. difficile* growth (Figure 1B). BHI contains 0.2% glucose as formulated and is unbuffered, likely accounting for the reduced pH. In conditioned SM with 0.4% added glucose and an average starting pH of 6.4, *C. difficile* growth was not supported (Figure 1D). These data suggest an inhibitory pH threshold of around 6 for *C. difficile* in VRE conditioned media. We note that *C. difficile* itself is capable of lowering medium pH when in monocoulture in both SM (pH 6.5) and BHI (pH 5.7) with added glucose, however, VRE lowers pH by an additional unit in both media (Figure S1). When fructose is added to co-cultures during stationary phase, *C. difficile* CFUs were significantly reduced along with a corresponding drop in pH (Figure S2). These data suggest that inhibition is not limited exponential growth phase. This effect can be bactericidal as resuspending late log-phase *C. difficile* in VRE-conditioned high glucose media for 2 hours resulted in a significant reduction in *C. difficile* (Figure S3). To further test if acidification is necessary for inhibition of *C. difficile* growth in VRE-conditioned media, we grew VRE to exhaustion in SM +/− 0.6% glucose with 100 mM of HEPES, MOPS or PIPES buffer (Figure S4). Buffering with MOPS resulted in a rise in the pH of the conditioned medium from 5.7 to 6.1, which resulted in a partial rescue of *C. difficile* growth. Taken together these data suggest that in glucose rich media VRE produces organic acids which lower pH and inhibit *C. difficile*.

The inhibition of *C. difficile* growth described above could be the result of an inhibitory factor produced by VRE or it could be due to nutrient limitation. To differentiate between these two mechanisms, we first generated VRE conditioned SM in a range of glucose concentrations. Then we created a dilution series of those conditioned media in sterile PBS. VRE conditioned media inhibits *C. difficile* growth with 0.4% and 0.6% total glucose concentration (Figure 2). When VRE conditioned media containing 0.4% glucose is diluted 1:2 in PBS, it no longer inhibits *C. difficile* growth. These data confirm that VRE inhibits *C. difficile* in a manner consistent with the production of organic acids and not via nutrient limitation.

**Figure 2:**
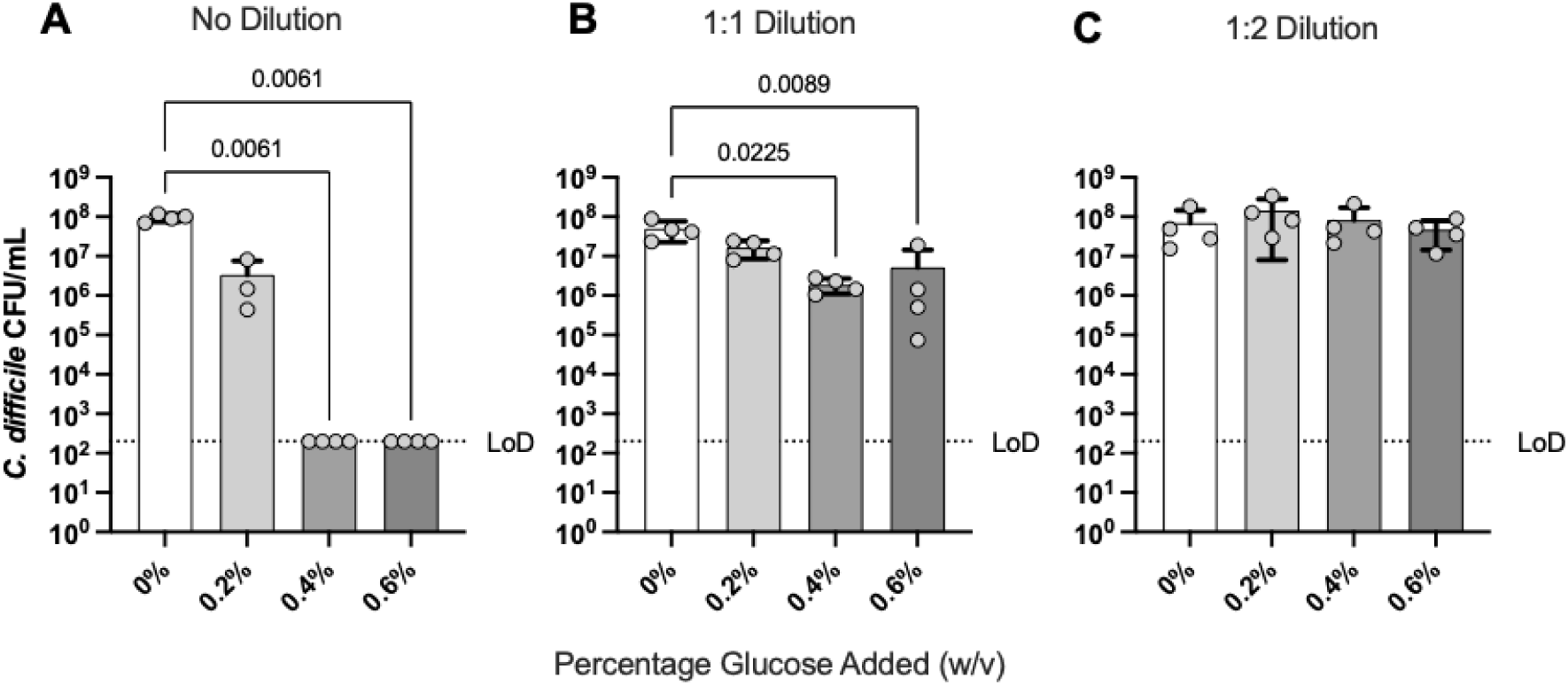
Inhibition by VRE-conditioned media is not caused by nutrient limitation. VRE conditioned sporulation media (SM) were generated with increasing concentrations of added glucose, filter sterilized, then kept undiluted (A) diluted 1:1 (B) or 1:2 (C) with reduced PBS before inoculation with *C. difficile* and plating at 48 hours of growth. Data are combined from 4 independent experiments (n=3-4 biological replicates per condition). Kruskal-Wallace 1-way ANOVA with Dunn’s correction vs. 0% glucose.

To determine if VRE-mediated acidification is necessary for inhibition of *C. difficile* we neutralized VRE conditioned BHI + 0.4% glucose (pH 5) with sodium hydroxide to pH 7. In the neutralized conditioned media *C. difficile* grew to similar levels as in naive BHI + 0.4% glucose (Figure 3A), suggesting that acidification is necessary for inhibition of *C. difficile* under these conditions. To determine if acidification of media is sufficient to inhibit *C. difficile* growth we acidified naive BHI and SM (pH 7.0) with hydrochloric acid and found that *C. difficile* growth was inhibited at a pH between 5.0-4.5 in acidified BHI and 6.0-5.5 in acidified SM (Figure 3B). Finally, to determine if acidification has the potential to synergize with other secreted effectors in VRE conditioned media, we used HCl to acidify VRE-conditioned SM, which had a starting pH of 7.6, to a range of pH from 7.0 to 4.5 Here we observed an inhibitory pH between 6.0-5.5, similar to the threshold observed in HCl acidification of naive SM (Figure 3C). These data suggest that acidification is the primary inhibitor of *C. difficile* under these in vitro conditions.

**Figure 3:**
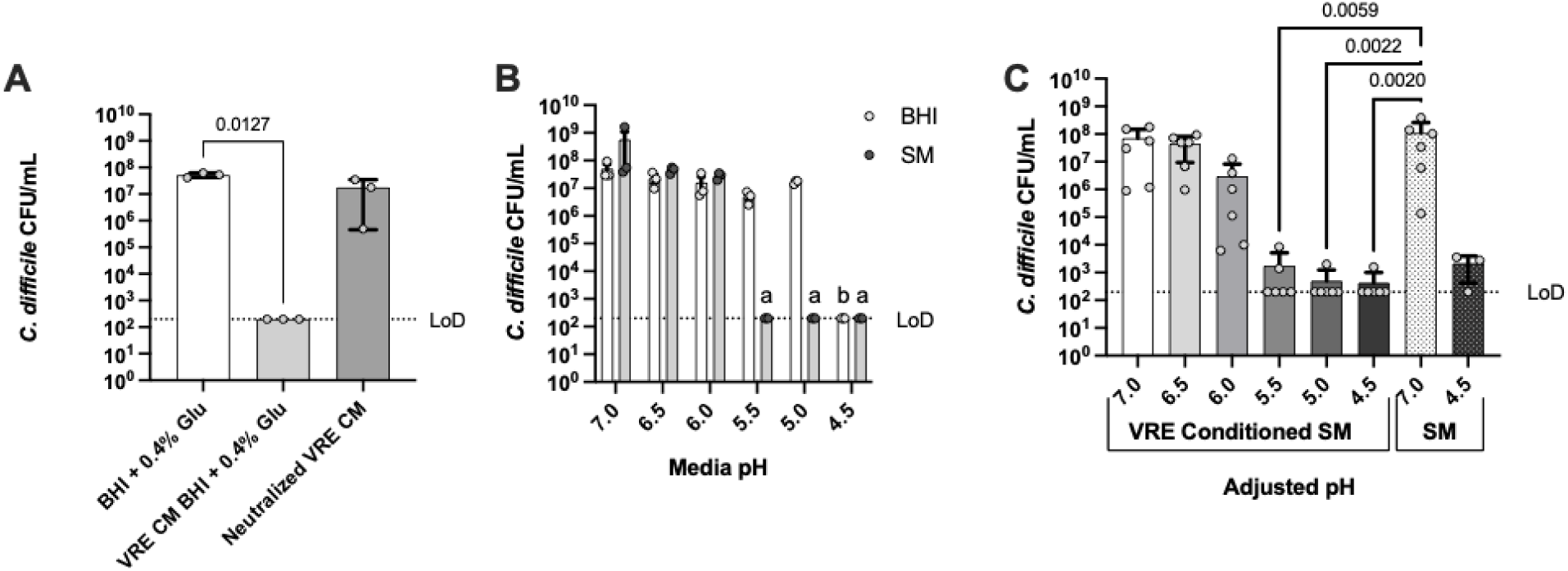
Acidification is necessary and sufficient for inhibition of *C. difficile*. A) VRE-conditioned BHI + 0.4% glucose or neutralized with NaOH to a pH of 7 and inoculated with *C. difficile* for 48 hours before plating and compared to *C. difficile* growth in naive medium at a pH of 7.0. Data are combined from 3 independent experiments, n=3 biological replicates, Kruskal-Wallis One-way ANOVA with Dunn’s correction vs. BHI + 0.4% Glu. B) BHI and SM were acidified with reduced HCl in 0.5 unit pH increments before inoculation with *C. difficile* and growth for 48 hours. Data are combined from 3 independent experiments, n=3 biological replicates, a: p = 0.0351, b: p = 0.0084, Kruskal-Wallis One-way ANOVA with Dunn’s correction versus pH 7.0 CFUs for each medium. C) VRE conditioned SM was filter-sterilized and acidified with reduced HCl before inoculation with *C. difficile* for 48 hours. Data are combined from 3 independent experiments, n=6 biological replicates, Kruskal-Wallis One-way ANOVA with Dunn’s correction versus pH 7.0 SM CFUs for each condition.

### VRE can metabolize specific carbon sources into organic acid to inhibit *C. difficile*

Next we tested the hypothesis that the acidification of any sugar is sufficient to cause VRE to inhibit the growth of *C. difficile* in conditioned media. We used a panel of simple sugars that differed in their reported metabolism by enterococci. Glucose and fructose were reported to be acidified, while fucose and xylose were reported not to be acidified (Schleifer & Kilpper-Balz, 1984). Tagatose and arabinose acidification was reported to differentiate between *E. faecalis* and *E. faecium* (Schleifer & Kilpper-Balz, 1984). We also included the disaccharide trehalose and the polysaccharide inulin which have been implicated experimentally in mouse models of *C. difficile* infection (Collins et al., 2018; Hryckowian et al., 2018). We generated VRE conditioned SM with 0.6% of each sugar and grew *C. difficile* for 48 hours before plating to simulate biofilm culture conditions. We found that only VRE conditioned SM containing glucose, fructose and trehalose inhibited growth of *C. difficile* when compared to growth in the same naive media (Figure 4A). Glucose, fructose and trehalose lowered pH of the conditioned media significantly with mean pH of 6.1, 6.0 and 5.7, respectively. These pH levels are consistent with the inhibitory pH levels described above.

**Figure 4:**
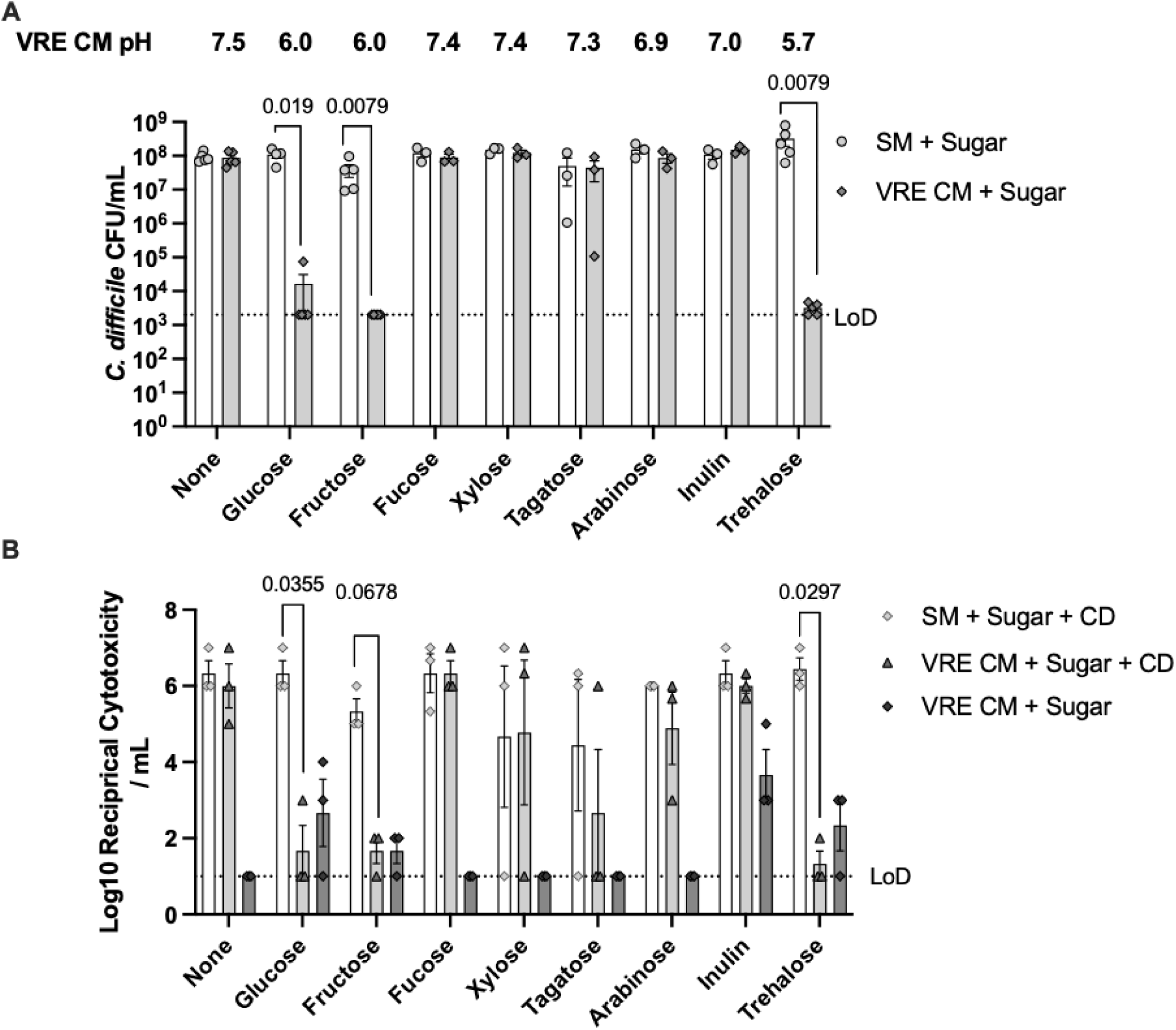
Inhibition of *C. difficile* by VRE is carbon source dependent. A) *C. difficile* was grown in fresh SM + 0.6% of the carbon source (SM + Sugar) indicated on the x-axis or VRE conditioned SM + 0.6% carbon source (VRE CM + Sugar). Starting mean pH of VRE CM + Sugar for each condition is reported above the graph. Data are combined from 3-5 independent experiments per condition, Mann-Whitney test. B) Cytotoxicity on Vero cells of filtered supernatants from A. Data are representative of 2 independent assays, Kruskal-Wallis One-way ANOVA with Dunn’s correction versus SM + Sugar + CD for each condition.

We performed cytotoxicity assays on Vero cells using filtered supernatants from the carbon source screen described above. We found significant reductions in cytotoxicity of VRE conditioned SM containing the acidified carbon sources, glucose, fructose or trehalose, which reflects the killing of *C. difficile* by VRE (Figure 4B). It is important to note that although glucose can suppress toxin production via catobolite repression and the transcription factor CcpA during exponential phase (Antunes et al., 2011; Dupuy & Sonenshein, 1998; Hofmann et al., 2021), we did not observe a decrease in cytotoxicity in samples containing *C. difficile* grown in naive SM + 0.6% glucose compared to SM with no carbon source. Our samples were collected at 48 hours post-inoculation, well beyond the exponential phase and likely reflects the total accumulation of toxin over 48 hours of growth. Additionally, in supernatants collected from VRE monocultures, we observed an unexpected but statistically significant increase in cytotoxicity from VRE conditioned SM + 0.6% inulin. This cytotoxicity was not neutralized by *C. difficile* anti-toxin.

To test the effects of carbon source availability in a more physiologically relevant in vitro system, we utilized extracts from cecal content of germ free mice and compared growth of *C. difficile* in naive and VRE conditioned filter-sterilized cecal extracts. In cecal content conditioned by VRE, we observed a slight, but not statistically significant (p = 0.13531) increase in *C. difficile* growth after 48 hours (Figure 5A) which agrees with recent findings that *E. faecalis* can promote *C. difficile* growth *in vivo* (Smith et al., 2022). However, when we added 0.6% glucose as a carbon source to the cecal extract before VRE conditioning, growth of *C. difficile* was not supported and pH was reduced to 4.01. Addition of 0.6% fucose, which is not acidified by VRE, as a carbon source did not result in a significant difference in *C. difficile* growth or pH (6.28) between naive and VRE conditioned cecal extract. These data suggest that the byproducts of carbon source metabolism by VRE can suppress *C. difficile* growth in cecal extracts. In these conditions we did not observe a significant reduction in cytotoxicity of VRE conditioned cecal content extract containing glucose, despite suppression of *C. difficile* growth below the level of detection (Figure 5B).

**Figure 5.**
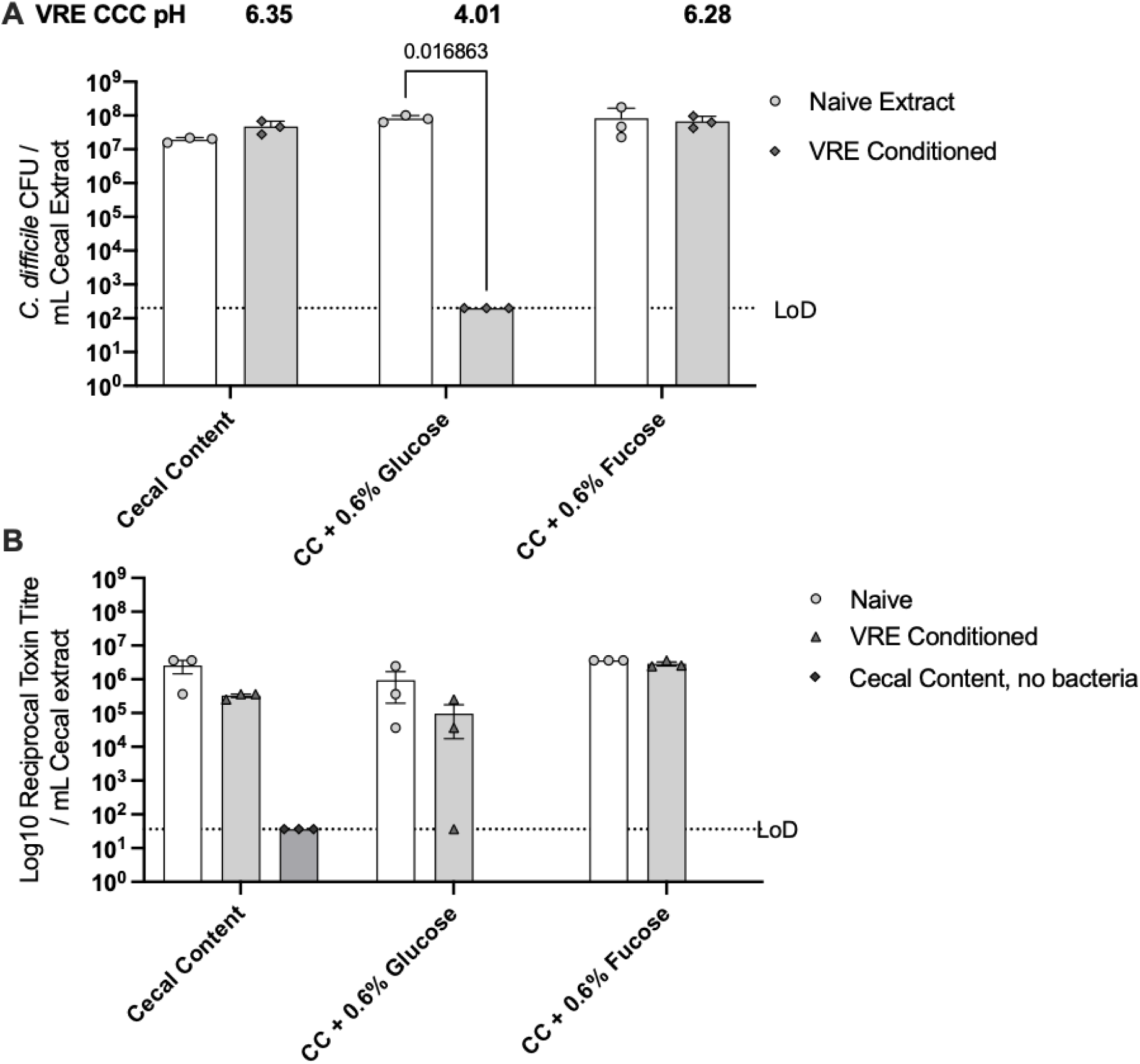
Acidification of glucose inhibits *C. difficile* growth in germ-free cecal content extracts. A) Sterile filtered extracts of cecal content from germ-free mice were prepared with reduced PBS and the indicated added carbon source or equivalent volume PBS and inoculated with VRE and grown for 24 hours. Conditioned cecal content (VRE CCC) were sterile filtered then inoculated with *C. difficile* and grown for 48 hours followed by plating on selective media. Data are combined from 3 independent experiments, n = 3 biological replicates. Unpaired Welch’s t-test. B) Vero cell toxin tire assay of filtered supernatants following *C. difficile* culture. Data are representative of two independent experiments, Unpaired Welch’s t-test.

We then tested if acidification-mediated suppression of *C. difficile* is conserved among enterococci. We generated conditioned SM + 0.6% glucose from a panel of enterococci including *E. hirae*, and vancomycin sensitive isolates of *E. faecuim* and *E. faecalis*. All of the tested strains completely suppressed *C. difficile* growth and produced conditioned media with a pH below 6 (Figure 6A). We then tested a small panel of *Clostridioides* species for growth in VRE conditioned media containing glucose (Figure 6B). Growth of *C. scindens, C. inoccuum, C. citrone, and C. difficile* ribotype 027 strain R20291 were inhibited below the limit of detection, while *C. bifermentans* showed significantly reduced growth. Acidification was necessary for growth inhibition of all strains as growth was restored when VRE conditioned media containing glucose was neutralized with NaOH. These data suggest that acidification is conserved among enterococci and is necessary and sufficient to inhibit growth of diverse *Clostridioides* species under these conditions.

**Figure 6:**
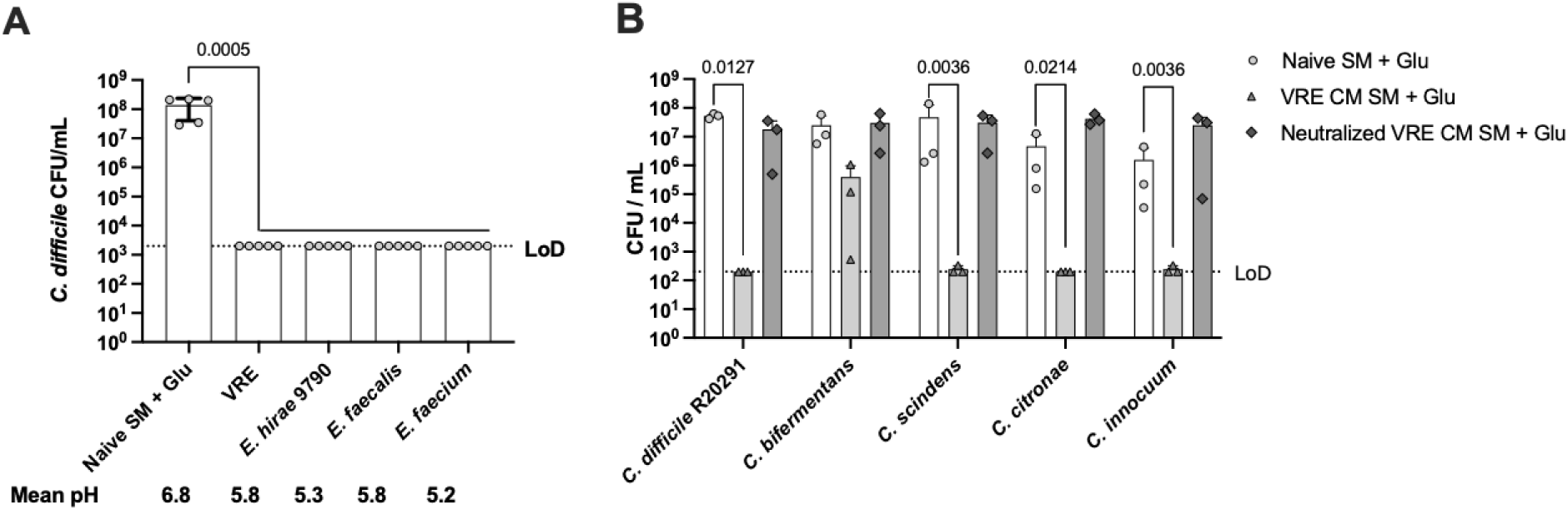
Acidification mediated inhibition is conserved among enterococci and clostridia. A) SM + 0.6 % glucose was conditioned by incubation with the indicated strains on the x-axis followed by inoculation with *C. difficile* and growth for 48 hours. Mean pH of the conditioned media are displayed below the x-axis (n=3). CFU data are combined from 3 independent experiments, n=5 biological replicates, Kruskal-Wallis One-way ANOVA with Dunn’s correction vs. Naïve SM + Glu B) VRE conditioned SM + 0.6% glucose was created by 48 hours of VRE growth followed by filter sterilization or was neutralized with NaOH to pH7 before filter sterilization and inoculation with *C. difficile* for 48 hours. Data are combined from 3 independent experiments, n=3 biological replicates, Kruskal-Wallis One-way ANOVA with Dunn’s correction vs. Naive SM + Glu.

### Fructose supplementation is not sufficient to restore colonization resistance in mice

To model the effects of supplementation with a fermentable carbon source, we adapted an existing mouse by supplementing with 15% w/v fructose in drinking water during *C. difficile* and VRE co-infection (Keith et al., 2020). Excess dietary fructose accumulates in the colon and alters the metabolism of VRE (Isaac et al., 2022; Jang et al., 2018). Mice were sensitized to infection through vancomycin and ampicillin in drinking water, followed by colonization with VRE which is resistant to both antibiotics (Keith et al., 2020). Fructose supplementation was added 24 hours before VRE challenge and maintained to the conclusion of the experiment (Figure 7A). Twenty-four hours after colonization with VRE, antibiotics were withdrawn for 48 hours followed by *C. difficile* infection for 24 hours followed by sampling of cecal content. Contrary to our hypothesis, supplementation with fructose in VRE colonized mice did not affect *C. difficile* colonization or toxin production (Figure 7B-C). There was a not significant trend towards higher fructose in fructose supplemented control mice (p = 0.07) when compared to naïve mice. However, we did not detect a significant change in fructose levels between CD + VRE mice and CD + VRE + Fructose. We also did not detect a significant change in pH of cecal content between dual colonized mice and dual colonized mice supplemented with fructose. We did, however, detect a rise in the mean pH between Naïve and antibiotics treated mice of 6.30 ± 0.02 to 6.85 ± 0.14 as has been reported in previously (Sorbara et al., 2019). These data suggest that the dietary supplementation attempted here was not sufficient to alter the pH of the cecum.

**Figure 7:**
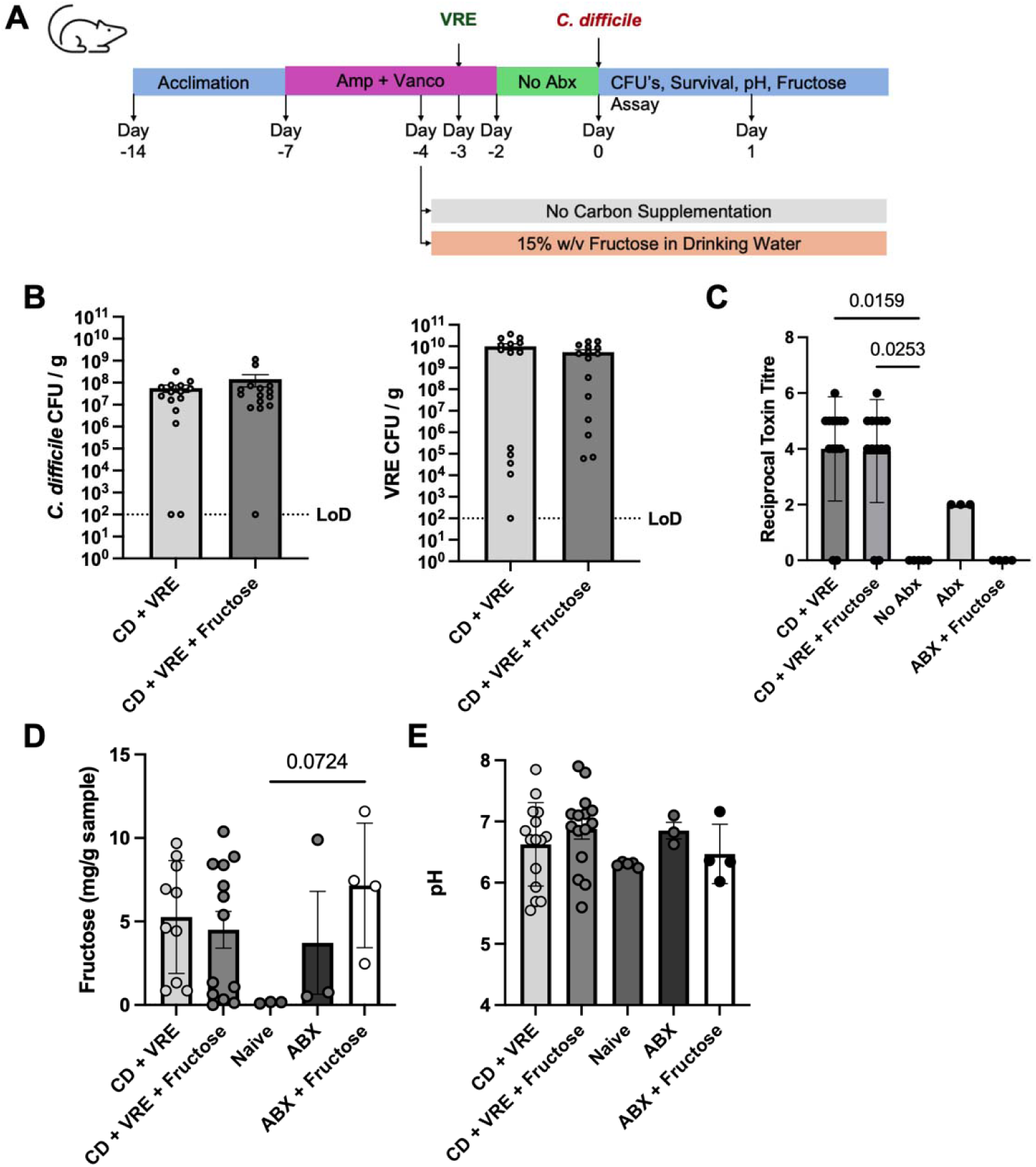
*C. difficile* – VRE dual-infection with fructose supplementation. A) Mice were treated with Ampicillin and Vancomycin in drinking water and 15% Fructose was added to the drinking water of one arm at day −4, followed by VRE colonization on day −3. Antibiotics were withdrawn for 48 hours followed by *C. difficile* infection on day 0. Data in B-E are combined from 3 independent experiments. B) *C. difficile* and VRE CFU levels in cecal content measured by selective plating. C) Vero Cell cytotoxicity of filtered cecal content extracts. D) Fructose levels in cecal content measured by an enzymatic assay. E) pH of cecal content. Data was collected with 4 −5 mice per condition combined from 3 independent experiments. Statistics: A and B, t-test with Welch’s correction. C-E, One-way ANOVA with Dunn’s correction.

## Discussion

We initially set out to develop an in vitro co-culture biofilm model of VRE and *C. difficile* using excess glucose to promote adherence of the bacteria to plastic plates (Ðapa et al., 2013). Growth of both species in excess glucose was not supported in co-culture (Figure 1). We found that acidification of glucose is necessary, sufficient and appears to be the primary mechanism by which VRE affects *C. difficile* growth in the presence of excess glucose in vitro (Figure 3). To test if VRE conditioned media inhibits *C. difficile* under all growth conditions we altered the available carbon source and found that growth inhibition is limited to contexts in which the carbon source is acidified by VRE. We observed a pH-mediated inhibition of growth in the presence of glucose, fructose and trehalose (Figure 4A). Added glucose also led to pH-mediated growth inhibition when the growth medium was germ-free mouse cecal content extract (Figure 5A). These data suggest that control of the available carbon source is critical to developing in vitro co-culture assays with lactic acid bacteria. We also measured the toxin titre of *C. difficile* in the carbon source screen and did not find significant changes in toxin production without an underlying inhibition of growth (Figure 4B). This is likely due to the 48-hour time point chosen for analysis, which is beyond the exponential phase in which toxin production is affected by carbon source in single species culture (Dupuy & Sonenshein, 1998). Given the importance of growth phase to toxin production, it is likely that competition for nutrients in dual-species culture may affect the induction of toxin at earlier time points (Smith et al., 2022).

We also detected what is potentially a novel toxicity of VRE towards cultured Vero cells. This cytotoxic activity was present when conditioned medium was prepared from SM containing 0.6% inulin, a fructan dietary fiber. The samples used in the assay were centrifuged and filtered with a 0.2 μm filter, suggesting that the cytotoxic substance is soluble and filterable. Inulin has generally enhanced protection in human fecal chemostat (Hopkins & Macfarlane, 2003) and mouse models of *C. difficile* infection (Hryckowian et al., 2018). If inulin induces cytotoxicity by VRE, it is possible that supplementation would result in increased virulence in co-infected mice.

In mouse models of infection, *C. difficile* first begins to accumulate in the caecum 6-12 hours post infection (Koenigsknecht et al., 2015). The colon, particularly the cecum, is densely colonized and is a site of high degrees of population diversity and countless metabolic products. A general feature of the mouse (Shimizu et al., 2021; Sorbara et al., 2019) and human cecum (Koziolek et al., 2015; Nugent et al., 2001) is an acidic pH of 5.0-6.5 at steady state (Brinck et al., 2025). During *in vitro* culture *C. difficile* growth and sporulation is limited following as little as a half-unit shift in pH (Wetzel & McBride, 2020). Growth, sporulation and germination are almost completely inhibited at pH of less than 6 (Kochan et al., 2018; Wetzel & McBride, 2020; Woo et al., 2011). Treatment with antibiotics in mice raises colon pH from acidic to slightly alkaline conditions, suggesting that the gut microbiota are necessary for maintaining acidic pH (Shimizu et al., 2021; Sorbara et al., 2019). These data suggest that the pH of the gut lumen, unlike the pH of blood, is not tightly buffered and may be among the many factors that contribute to *C. difficile* colonization resistance mediated by the gut microbiota. Most bacteria are capable of living in a pH range of 3-4 units and pH may act as a constraint on growth of commensals and opportunistic pathogens (Jin & Kirk, 2018). We found that pH-mediated inibition of *C. difficle* is at least partly bactericidal (Figure S3). Given that ingested *C. difficile* spores germinate well in the pH neutral small intestine (Koenigsknecht et al., 2015), it is possible that transit into the acidic colon (pH ∼ 6) is a natural restraint on growth and contributes to colonization resistance. In humans, a small study found a significant association between an alkaline fecal pH and *C. difficile* infection (Gupta et al., 2016).

The pH of the gut lumen is modulated and constrained by inputs from the host diet, and the metabolism of the gut microbiota and the colonic epithelium. The colonic epithelium produces and secretes bicarbonate, which increases the alkalinity of the gut lumen (Alka & Casey, 2014), which could potentially buffer acid production by VRE in our model. However, several studies have noted decreased levels of the bicarbonate transporter DRA (SLC26A3) in the colon of *C. difficile* infected mice (Coffing et al., 2018; Peritore-Galve et al., 2023; J. Wang et al., 2020). In addition to DRA, toxin-dependent damage also reduced the expression of the host glucose transporter SGLT1 in mice and resulted in an increase in the level of glucose in the stool (Peritore-Galve et al., 2023). The colonic epithelium also produces lactic acid as a byproduct of glycolysis which can be converted into short chain fatty acids by the gut microbiota (Louis et al., 2022). On the bacteria side of this interaction, in cells of *C. difficile* during active infection, toxin-mediated damage causes the upregulation of carbohydrate PTS importers (Fletcher et al., 2021). Carbohydrate metabolism transcripts as a class, including genes involved in fructose metabolism were increased by both *C. difficile* and *E. faecalis* OG1RF during co-culture in vitro. Taken together, these data suggest that *C. difficile* and VRE may import and metabolize excess fructose in the colon. Primary metabolism by the gut microbiota produces weak organic acids such as lactic acid and short chain fatty acids when grown in a fermentable carbon source (Gregory et al., 2021). The short chain fatty acid butyrate is growth suppressive to *C. difficile* in vitro and is inversely correlated with colonization in mice and humans (Pensinger et al., 2024, 2023). All these factors together have the capacity to influence the standing pH of the large intestine.

In certain cases, such as in critically ill patients treated with antibiotics who become dominated by VRE (Liao et al., 2021), it may be desirable to enhance the acidity of the gut lumen through administration of a diet or prebiotics that results in acid production and suppression of *C. difficile*. Our data suggest that supplementation with fructose in drinking water alone was not sufficient to affect *C. difficile* colonization in VRE co-infected mice, nor was it sufficient to affect the pH of the contents of the cecal lumen (Figure 7). It is possible that more complex formulations of dietary input (Hryckowian et al., 2018; Mefferd et al., 2020) may be necessary to restore *C. difficile* colonization resistance in a low diversity gut microbiota.

Clinical trials have tested the efficacy of probiotic lactic acid bacteria on *C. difficile* infections with mixed results (Goldstein et al., 2017; Mills et al., 2018). It is clear, however, that strains from genus *Bifidobacterium* and *Lactobacillus* can inhibit *C. difficile* in vitro by decreasing pH (Fredua-Agyeman et al., 2017) and in some cases similar probiotics have reduced *C. difficile* virulence in animal models of infection (Wei et al., 2018). For example, lactic acid production by *Streptococcus thermophilus* lowered the pH of conditioned media and was inversely correlated with *C. difficile* growth and toxin production *in vitro* and in a mouse model of infection (Kolling et al., 2012). These data suggest that acidification-mediated effects on growth of *C. difficile* are not limited to enterococci and may be a general feature of lactic acid bacteria and any bacteria capable of acidifying a carbon source.

Multiple groups have reported that changes in pH affect growth and survival of core taxa of the gut microbiota. For example, during in vitro culture at pH 5.5 multiple human gut isolates from genus *Bacteroides* failed to grow (Duncan et al., 2009). This pH sensitivity of *Bacteroides* has been replicated in several batch culture fermenter studies seeded with donor human feces (Firrman et al., 2022; Ilhan et al., 2017; S. P. Wang et al., 2020). These studies also consistently reported reductions in levels of genus *Clostridioides* (or *Clostridium*) under low pH conditions. In keeping with the batch fermentation studies mentioned above, we also found that commensal *Clostridioides* species are subject to a similar pH-mediated sensitivity to VRE-conditioned media as *C. difficile* (Figure 6). Therefore, efforts to remediate dysbiosis in recurrent *C. difficile* infection by reintroducing commensal spore-forming anaerobes (Feuerstadt et al., 2022; Louie et al., 2023) may benefit from a different strategy. Here, temporarily limiting acid production by enterococci and other lactic acid bacteria through dietary intervention may favor engraftment of the bacteriotherapy. The conundrum here is similar to that faced by chemists since the beginning of antibiotics development: How do you specifically target pathogenic bacteria, while sparing or promoting the closely related commensal strains necessary for steady-state health? A solution will require a more complete knowledge of the metabolism of pathogenic and commensal clostridia.

## Acknowledgements

Work in this study was supported by startup funds from Binghamton University, the SUNY Research Foundation and NIAID R21AI171634. PTM is an inventor on US Patents 10,646,520, 11,207,374 and 11,471,495 owned by Memorial Sloan Kettering Cancer Center and receives licensing royalties originating from Seres Therapeutics, Inc. and Nestle Health Sciences. We thank Eric Pamer, Alexander Rudensky, John Hambor and Su-Ellen Brown for bacterial strains. Germ free mouse cecal content was acquired using funds from Boehringer Ingelheim Pharmaceuticals, Inc. for an unrelated project.

---

**Figure S1:**
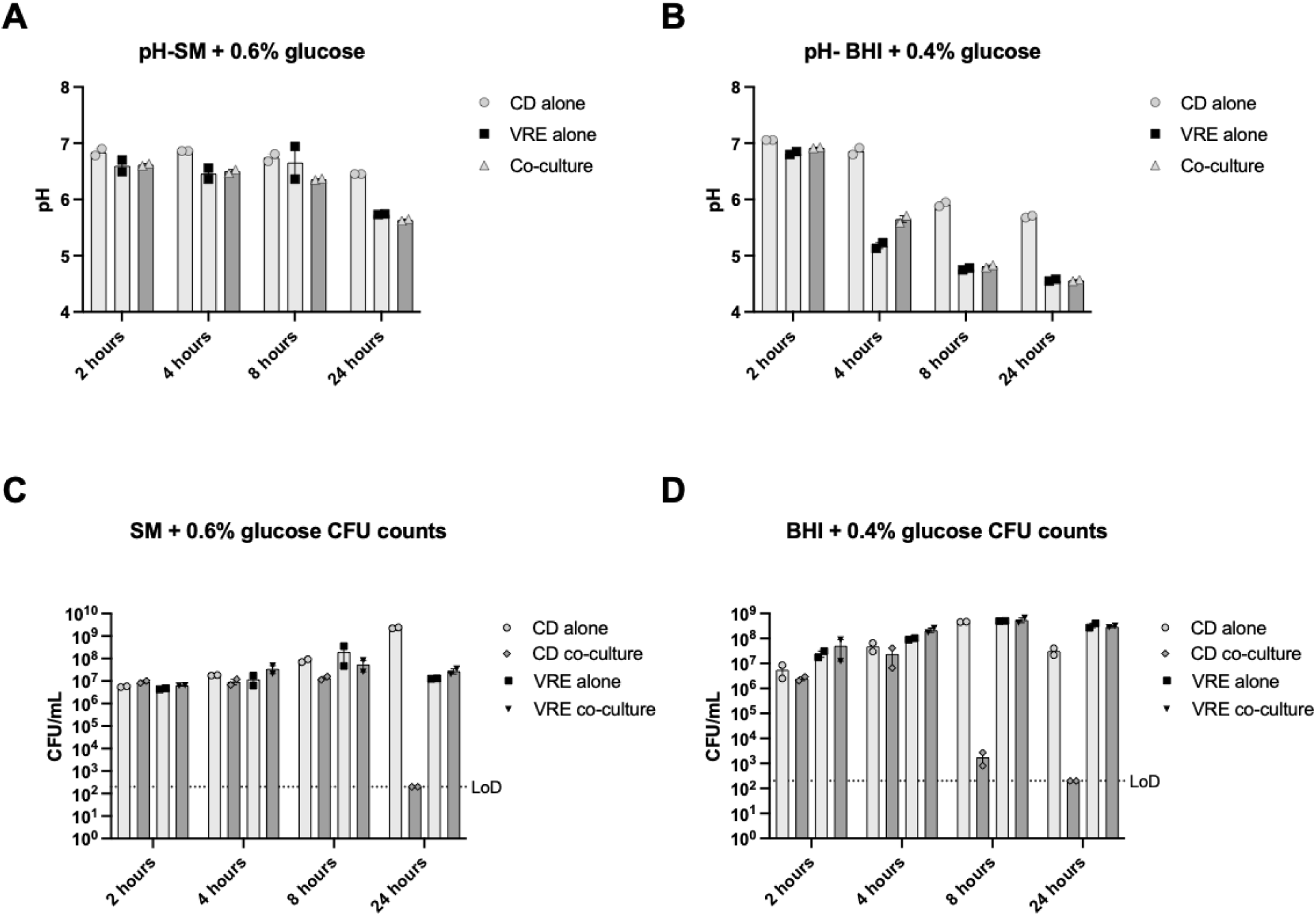
Dynamics of suppression of *C. difficile* by VRE. *C. difficile* and VRE were inoculated into SM + 0.6% glucose and BHI + 0.4% glucose. Samples were taken at 2 hours, 4 hours, 8 hours and 24 hours for pH and selective plating for CFU’s. A) pH of time course in SM + 0.6% glucose. B) pH of time course in BHI + 0.4% glucose. C) CFU’s corresponding to part A pH values. D) CFU’s corresponding to part B pH values. Data are combined from two experiments one biological replicate each.

**Figure S2:**
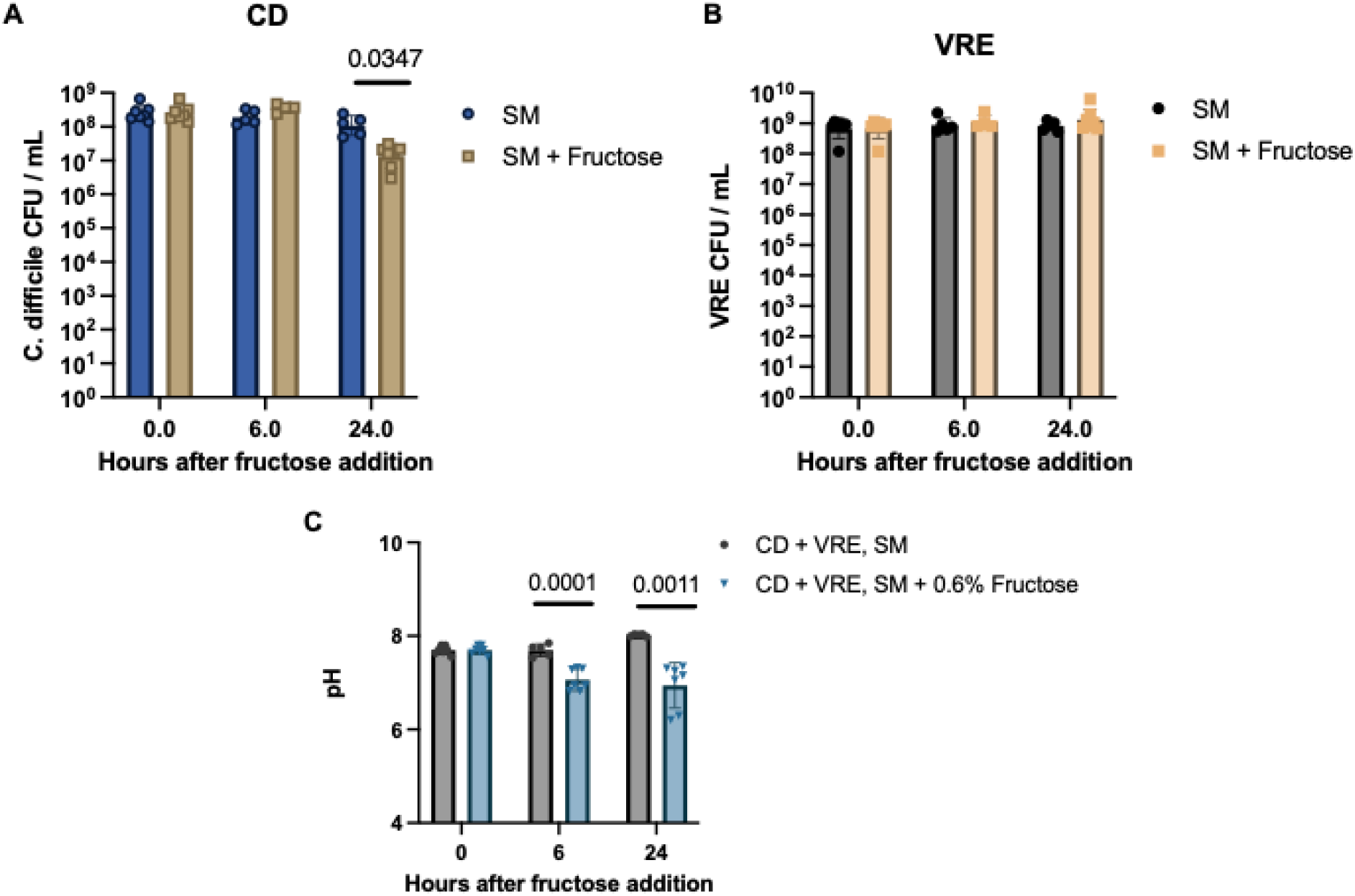
Addition of fructose at stationary phase is sufficient to inhibit *C. difficile* in co-culture. *C. difficile* - VRE co-culture was grown to stationary phase and 0.6% fructose or an equivalent volume of sterile water was added to the medium. Cultures were grown for the indicated time, plated to quantify CFUs and the pH of the medium was measured. A) *C. difficile* CFUs. B) VRE CFUs. C) pH of the medium at the indicated time points. Data are combined from 3 independent experiments with 4-7 biological replicates per time point. Statistics: Welch’s unpaired t-test

**Figure S3:**
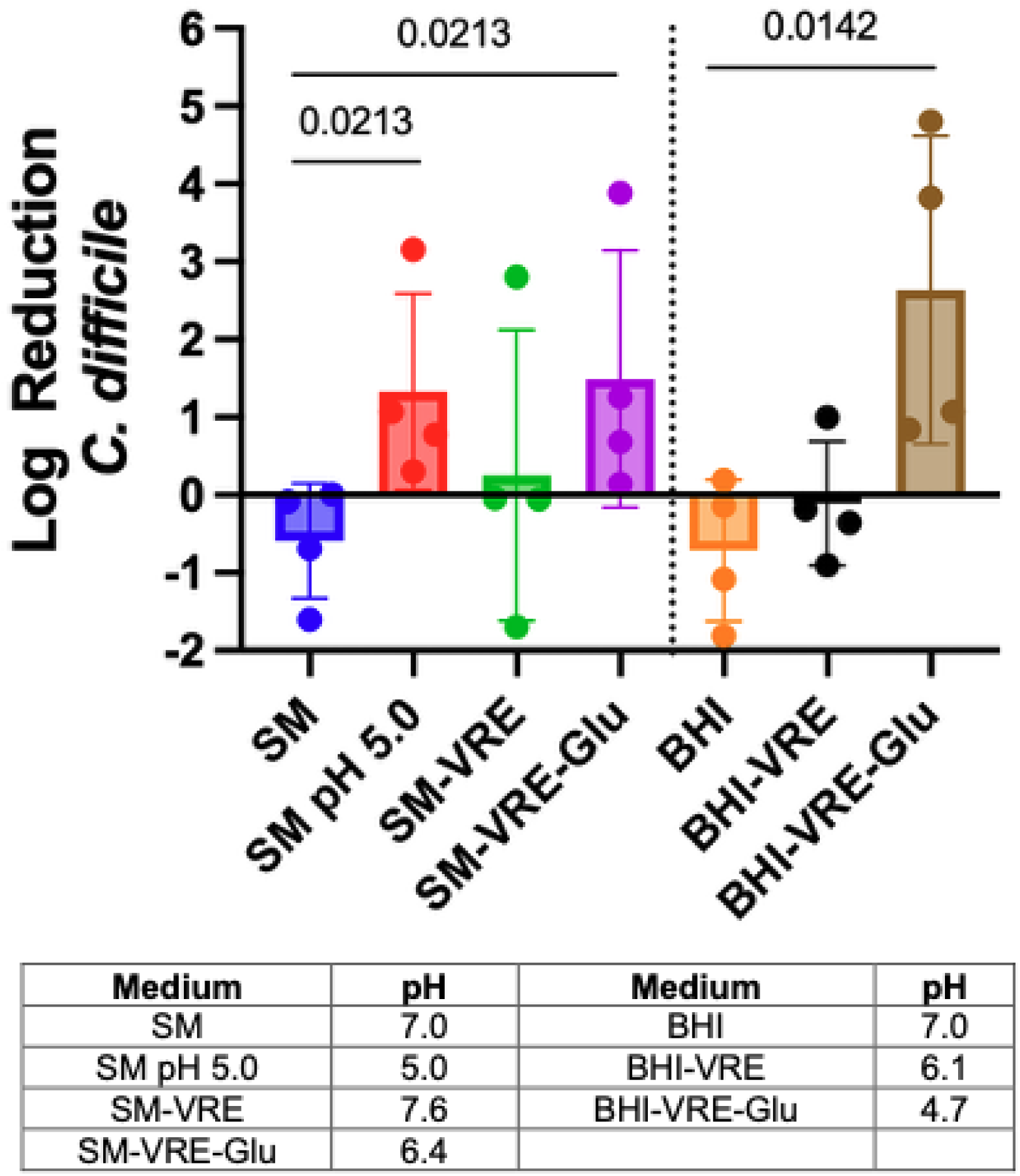
*C. difficile* inhibition driven by high glucose VRE conditioned media is at least partially bactericidal. *C. difficile* was grown to late log phase then pelleted and resuspended in: SM: Naïve SM, SM pH 5.0: SM pH-adjusted to 5.0 with HCl, SM-VRE: VRE conditioned SM, SM-VRE-Glu: VRE conditioned SM + 0.6% Glucose, BHI: Naïve BHI, BHI-VRE: VRE conditioned BHI, BHI-VRE-Glu: VRE conditioned BHI + 0.4% Glucose. After 2 hours of incubation at 37C, *C. difficile* was quantified by selective plating. Data are shown as log reduction compared to *C. difficile* resuspended in PBS. Table contains average pH values from 4 experiments. Statistics Kruskal-Wallace One-way ANOVA versus naïve medium, n = 4 biological replicates.

**Supplementary Figure 4:**
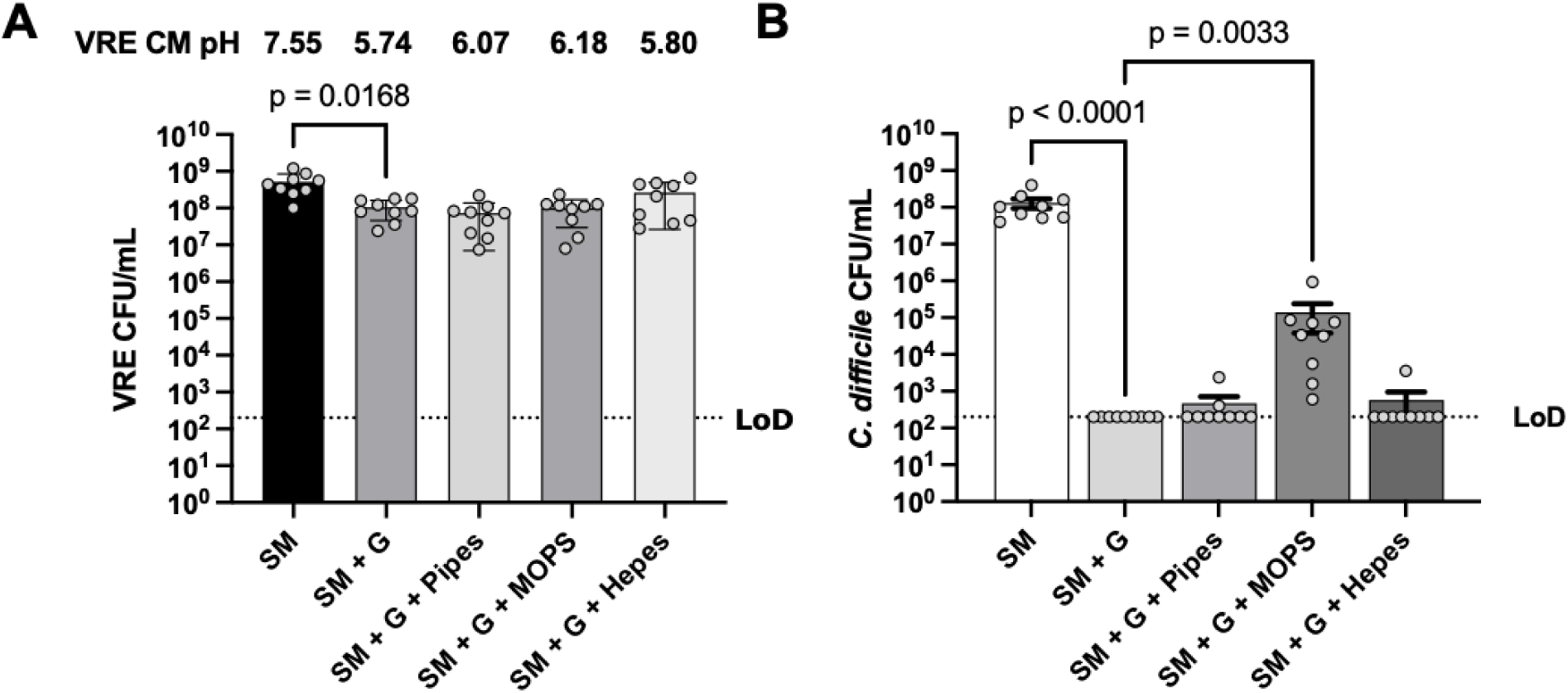
Buffering partially restores *C. difficile* growth in high glucose conditioned media. A) VRE was grown for 24 hours in SM liquid medium with and without 100mM buffers as labeled on the x-axis (G = 0.6% glucose), and plated for VRE. Medium was filter sterilized, pH was measured and used as growth medium for *C. difficile* in Figure S1B. Data are combined from 3 independent experiments, n = 9 biological replicates, Kruskal-Wallace one-way ANOVA with Dunn’s correction versus SM + G B) *C. difficile* CFU after 48 hours growth in VRE-conditioned medium, Kruskal-Wallace one-way ANOVA with Dunn’s correction versus SM + G.

**Table S1:**
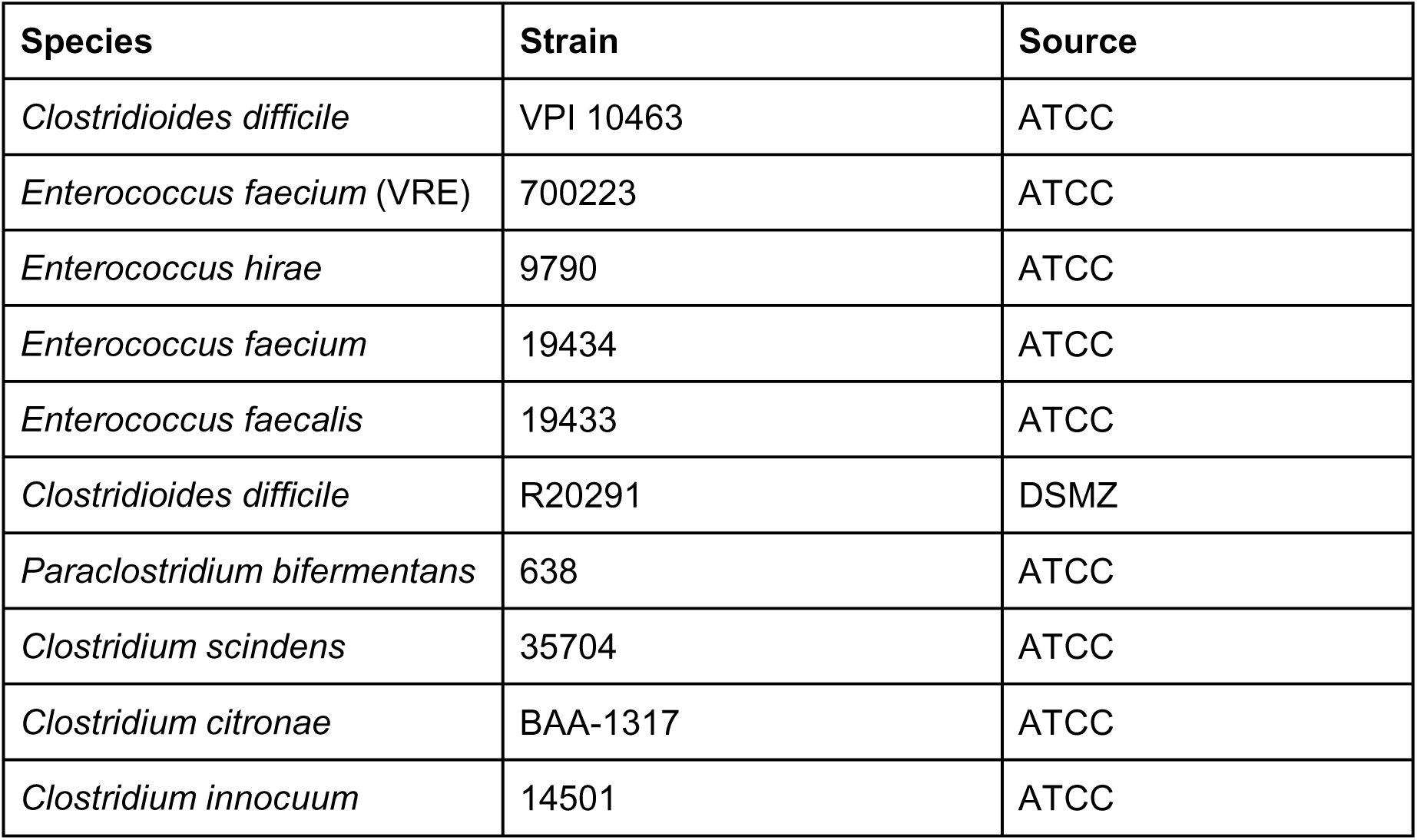
Table of Strains.

## Methods

### Strains and Growth Conditions

Strains are listed in Table 1. All experiments were conducted in an anaerobic chamber (Coy Laboratory Products), with an atmosphere of 90% N_2_, 5% CO_2_ and 5% H_2_. *C. difficile* and other clostridia strains were routinely cultured on BHI plates and liquid media (Brain Heart Infusion + 0.5% yeast extract) supplemented with 0.1% taurocholate and 0.3% L-cysteine. VRE and other enterococci were routinely cultured in BHI liquid medium and Enterococcosel (BD Biosciences) plates. For differential selection during co-culture, VRE and enterococci were plated on Enterococcosel (containing 8 μg / mL vancomycin and 100 μg / mL ampicillin for VRE) while *C. difficile* was plated on BHI plates supplemented with 0.1% taurocholate, 0.3% L-cysteine, 250 μg/mL D-cycloserine, 16 μg/mL cefoxitin. Commensal *Clostridia* were enumerated by plating on Columbia Blood Agar and counting by distinguishing between colony morphology of VRE.

### Media

BHI was made with 36g/L Bacto BHI powder with 5g/L yeast extract, bacterial, autoclaved, then 3 mL filter sterilized 10% L-cysteine was added to the media before aliquoting and reducing in the anaerobic chamber. Sporulation medium was made with 90g Bacto peptone, 5 g Bacto protease peptone, 1 g ammonium sulfate and 1.5 g tris base, then 3 mL 10% L-cystine was added after autoclaving. All liquid and plate media were pH adjusted to 7.0 with HCl unless otherwise specified before autoclaving. Glucose and other sugars used were dissolved into water at a 6% weight per volume ratio before being filter sterilized. Sugars were then added into the media before being placed in the anaerobic chamber or allowed to reduce in the chamber before being added to the media. Inulin was added as described with heating to allow full solubility before being filtered. All plates were poured in 18mL volumes and kept at 4°C until needed. 8-24 hours before using, plates were placed in the anaerobic chamber to reduce fully.

### *C. difficile* growth in VRE and enterococci conditioned media

Mid-log VRE overnight culture was OD-normalized to 0.5 before being inoculated (1:20 dilution) into SM + 0.6% sugar. The culture was grown for at least 24 hours anaerobically at 37°C until the OD_600_ was > 2 to ensure growth to saturation. Cultures were spun at 1500g x 15 minutes before being filter sterilized with a 0.22 μm filter syringe into 3 mL aliquots into glass tubes. 1.5 mL aliquots were also frozen at −80C for further use in cytotoxicity assays. *C. difficile* was inoculated 1:60 from mid-log into the VRE-conditioned media and incubated at 37°C for 48 hours before being serially diluted and drop plated for CFU/mL enumeration.

### Glucose titration in BHI and SM

BHI and SM were used to create media containing the following glucose concentrations: 0.0%, 0.2%, 0.4% and 0.6%. Media was placed in the anaerobic chamber to reduce before using. VRE was inoculated 1:20 into each glucose concentration of both BHI and SM. Cultures were incubated for 24 hours before being filtered sterilized as described above. *C. difficile* was inoculated 1:60 into VRE-conditioned media and fresh BHI and SM conditions and incubated for 24 hours before serial dilutions and drop plating for enumeration.

### Dilution of inhibition by VRE conditioned Media

VRE conditioned media was prepared by inoculating VRE 1:20 into BHI with 0.0%, 0.2%, 0.4% and 0.6% added glucose. Cultures were grown for 24-36 hours to an OD_600_ >2 before being spun down and filter sterilized. A subset of the supernatant was taken out of the chamber for pH recording. The VRE-conditioned media was then diluted 0, 1:1, 1:2, 1:4 and 1:8 with sterile PBS. After dilution, 3 mL aliquots were divided into test tubes per each biological replicate. *C. difficile* mid-log cultures were diluted 1:10 in sporulation media, then added to the aliquots in a 1:60 dilution and grown between 24 hour before drop plating for enumeration.

### Acidification and neutralization of SM and BHI

Acidification of SM and BHI was created by titrating HCl into the media to a pH of 7.0, 6.5, 6.0, 5.5, 5.0 and 4.5 before being autoclaved and aliquoted. *C. difficile* was inoculated into the reduced media 1:60 and incubated at 37°C for 48 hours before being serial diluted and drop plated for enumeration.

Neutralization was performed by growing VRE in SM + 0.6% glucose or BHI + 0.4% glucose to exhaustion before being spun down and filter sterilized. The filtered media pH was recorded to ensure full acidification at or below 5.5, then neutralized to a pH of 7.0 with NaOH. Neutralized media was filter sterilized again and aliquoted into 3 mL test tubes then placed in an anaerobic incubator for 24 hours to ensure full media reduction. 3 mL aliquots of the acidified media were collected to be used as a negative control. After being reduced, *C. difficile* was inoculated 1:60 into the media conditions. *C. difficile* was also inoculated 1:60 into fresh media at a pH of 7.0 as a positive control. All media conditions were incubated at 37°C for 48 hours before drop plating and enumeration.

### Cecal extract ex-vivo culture

Cecal content from adult germ-free C57BL/6 mice, 12-16 weeks old, were harvested under sterile conditions and frozen at −80C until use. Mice were maintained at an AAALAC accredited facility under an animal protocol approved by the Institutional Animal Care and Use Committee of Boehringer-Ingelheim Pharmaceuticals Inc. Cecal content extracts were prepared at 0.1g/mL of wet weight in reduced PBS in an anaerobic chamber. Cecal content extracts were prepared at 0.1g/mL of wet weight in reduced PBS in an anaerobic chamber. Contents were vortexed and then centrifuged at 1500g for 15 minute before filter sterilization with a 0.2 μm filter. The extract was split into three conditions with the following supplementation: 0.6% of PBS, glucose or fucose. VRE was inoculated 1:20 into half of the conditions described, grown for 24 hours, spun down and filter sterilized to create VRE conditioned extract. Mid-log *C. difficile* was inoculated 1:60 from an inoculum into naïve extract or VRE conditioned extract and grown for 24 hours at 37°C before serial dilution and drop plating.

### Cytotoxicity assays

Vero cells were grown at 10,000 cells per well in a 96-well plate in Eagle’s minimum essential media (EMEM) + 10% Fetal Bovine Serum and 1x Penicillin & Streptomycin overnight. Culture supernatants and mice samples were frozen at −80C until assay. All samples were spun down for 15 minutes at 15,000g before being filter sterilized into new tubes. Each sample was serial diluted down to 10^−6^ in a fresh 96-well plate with PBS. *C. difficile* purified toxin (TechLab) was diluted into 1 mL of sterile water as a positive toxin control. 96 µL of each sample was placed in the top row of a 96-well plate and 4 µL of the anti-toxin (TechLab) was added and incubated for at least 20 minutes to ensure full toxin neutralization from the anti-toxin. Samples with antitoxin were serially diluted in the remaining rows down to 10^−6^ in PBS. Once all toxin, aliquoted samples and antitoxin samples were completed, 100uL of each sample and the dilutions were placed onto their respective Vero-cell wells. Vero cells were incubated at 37°C with 5.0% CO_2_ overnight. Cytotoxicity was then analyzed with microscopy. Cells that showed rounding were positive for cytotoxicity. To calculate toxin titre, dilutions with less than 80% cell rounding were considered negative, and the previous dilution in the series was considered positive. This dilution number was then used to calculate the Log10 reciprocal toxin titre.

### Conservation of enterococci inhibition of clostridia through acidification

We first grew each enterococcal in BHI+0.6% glucose overnight, filter sterilized and inoculated *C. difficile* to grow for 48 hours at 37°C before serial dilution and drop plating on *C. difficile* plates for enumeration. We then used VRE to condition SM + 0.6% glucose, spun down and filter sterilized as previously described. We then inoculated mid-log cultures of the remaining clostridia panel into the VRE-conditioned media in a 1:60 dilution and grew *C. difficile* for 48 hours at 37°C before drop plating. We also performed neutralization and acidification as described above with this clostridia panel.

### Buffering of VRE CM in SM + 0.6% Glucose

VRE was grown overnight to an OD of >1.8 in SM or SM + 0.6% glucose with the following buffers; 100mM PIPES, 100mM HEPES, 100mM MOPS. After VRE growth, the samples were plated on enterococcus selective media, the samples were spun down and filter sterilized. An aliquot from each condition was taken to record pH. *C. difficile* was inoculated 1:10 into each VRE conditioned media and a naive set of conditions. *C. difficile* was grown for 48 hours before selectively plating.

### Mouse co-infection

C57BL6 mice, 7 weeks old, were purchased from Jackson Laboratories and were maintained under an approved protocol of the Binghamton University Institutional Animal Care and Use Committee (21-852). Mice were screened for *C. difficile* upon receipt by enrichment culture in CC-BHIS-TA medium (Maslanka et al. 2020). Mice were acclimatized for 1 week prior to antibiotics treatment and were maintained in sterile disposable individually ventilated caging with sterile bedding and irradiated food (LabDiet Rodent Diet 20). Mice were treated with 500 mg/L Ampicillin + 250 mg/L vancomycin in drinking water for 4 days, switched to water without antibiotics for 48 hours followed by infection with 2500 CFU of *C. difficile* VPI-10463 by oral gavage. Mice receiving VRE were gavaged with VRE after 3 days of antibiotics treatment. For mice receiving dietary supplementation, 15% fructose was added to drinking water after 3 days of antibiotics treatment. Samples were collected and transferred immediately into a pre-reduced anaerobic jar (AnareoPack-Anaero) to minimize oxidative damage to vegetative *C. difficile* during sampling and transport.

### Enumerating bacteria of *in vivo* models

Fecal pellets were collected from mice into sterile, pre-weighed microcentrifuge tubes on the day of acclimation, the day of VRE challenge, the day of *C. difficile* challenge and 24 hours after. VRE and *C. difficile* screening during acclimation was done using selective plating. Commensal enterococci was plated on non-selective enterococci plates and was maintained through the duration of the experiments. 24 hours after *C. difficile* challenge, mice were euthanized using an Euthanix lid for 10 minutes before cervical dislocation. Cecal content was collected into two tubes, one for CFU plating and one for pH measurements. To calculate CFU’s per 1g sample, tubes were weighed before and after sample collections and the raw CFU’s were divided by the sample weight. During *C. difficile* infection, tubes were placed in an anaerobic box before collecting. Samples were quickly collected from each mouse and put back into the anaerobic box before processing. All samples were resuspended in 1.0 mL pre-reduced PBS before plating. To enumerate spores, samples were diluted 1:1 in PBS and heat treated at 60°C for 30 minutes and vortexed at 15 minutes. Samples were frozen at −80°C until further use after initial CFU plating. Cecal pH was collected by dissolving cecal contents in 200uL nanopure water before pH measurement using an Accumet micro-electrode (Fisher).

### Fructose Assay

Cecal contents used for CFU plating and cytotoxicity assays were used for fructose concentration analysis by a colorimetric assay kit (NovusBio). The protocol was followed with the following adaptions: a standard curve was created using 0, 500, 800, 1000, 1600, 1800 and 2000 µg/mL fructose and the linear equation was calculated of y= 0.0002x + 0.0831. 25 uL of sample was combined with 1.5mL assay kit solution and boiled for 8 minutes at 100C. A Nanodrop (Thermo-Fisher) was blanked with nanopure water and measured at 285nm to analyze standard values and all sample values.

