## Supplemental Figures S1-S4 Table S1 for "Acidification-dependent suppression of *C. difficile* by pathogenic and commensal enterococci"

**Supplementary Figure S1**

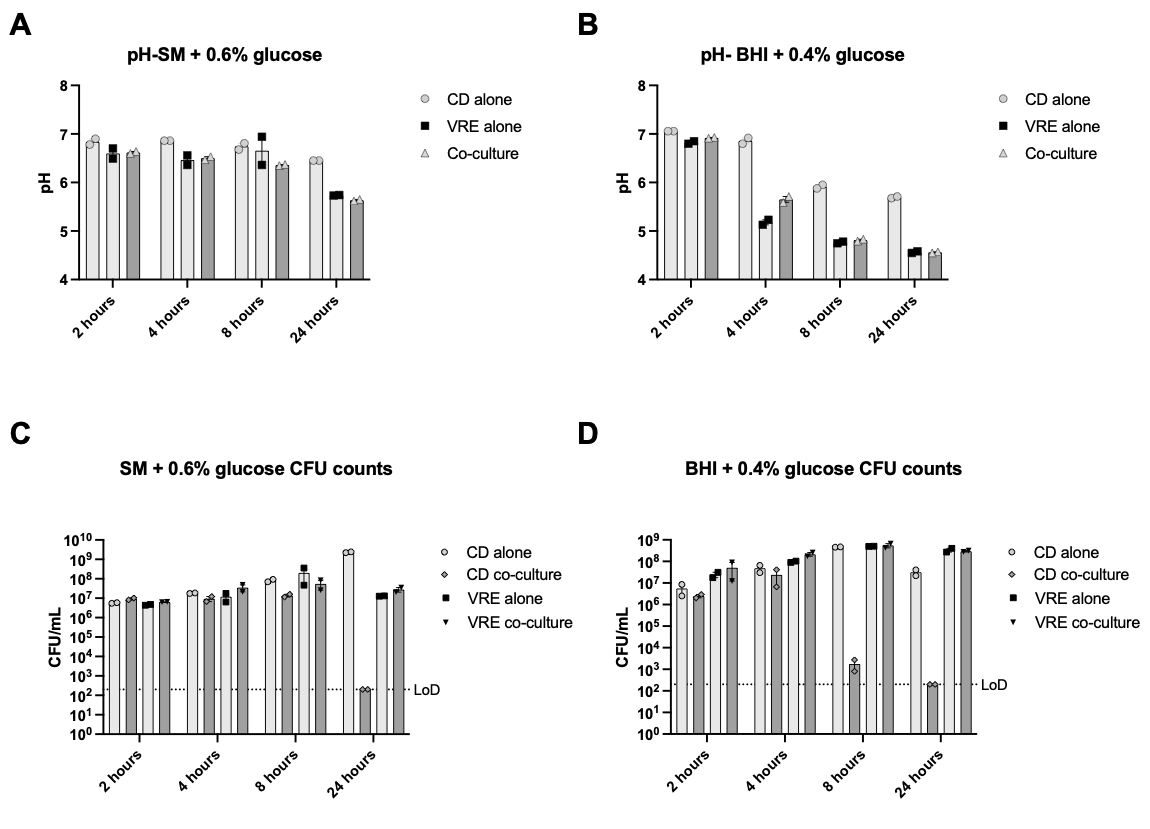

**Figure S1: Dynamics of suppression of *C. difficile* by VRE.** *C. difficile* and VRE were inoculated into SM + 0.6% glucose and BHI + 0.4% glucose. Samples were taken at 2 hours, 4 hours, 8 hours and 24 hours for pH and selective plating for CFU's. A) pH of time course in SM + 0.6% glucose. B) pH of time course in BHI + 0.4% glucose. C) CFU's corresponding to part A pH values. D) CFU's corresponding to part B pH values. Data are combined from two experiments one biological replicate each.

**Supplementary Figure 2**

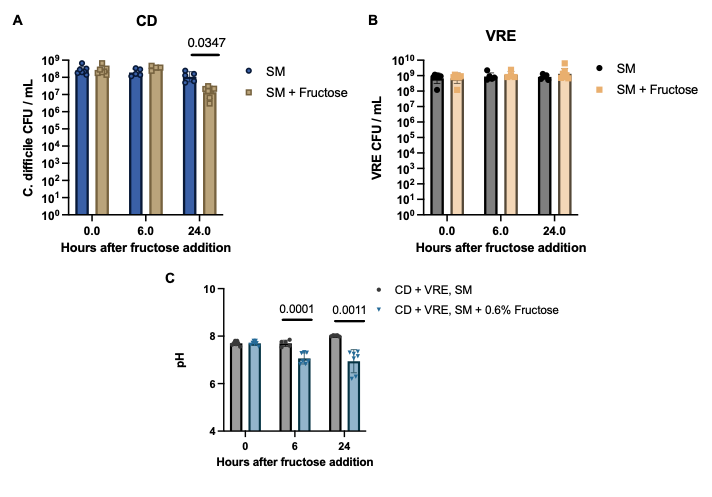

**Figure S2: Addition of fructose at stationary phase is sufficient to inhibit *C. difficile* in co-culture.** *C. difficile* - VRE co-culture was grown to stationary phase and 0.6% fructose or an equivalent volume of sterile water was added to the medium. Cultures were grown for the indicated time, plated to quantify CFUs and the pH of the medium was measured. A) *C. difficile* CFUs. B) VRE CFUs. C) pH of the medium at the indicated time points. Data are combined from 3 independent experiments with 4-7 biological replicates per time point. Statistics: Welch’s unpaired t-test

**Supplementary Figure S3**

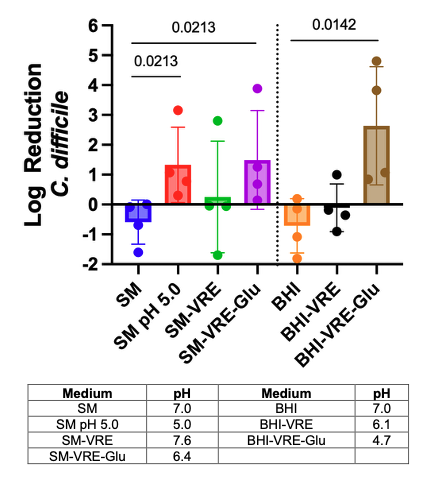

**Figure S3: *C. difficile* inhibition driven by high glucose VRE conditioned media is at least partially bactericidal.** *C. difficile* was grown to late log phase then pelleted and resuspended in: SM: Naïve SM, SM pH 5.0: SM pH-adjusted to 5.0 with HCl, SM-VRE: VRE conditioned SM, SM-VRE-Glu: VRE conditioned SM + 0.6% Glucose, BHI: Naïve BHI, BHI-VRE: VRE conditioned BHI, BHI-VRE-Glu: VRE conditioned BHI + 0.4% Glucose. After 2 hours of incubation at 37C, *C. difficile* was quantified by selective plating. Data are shown as log reduction compared to *C. difficile* resuspended in PBS. Table contains average pH values from 4 experiments. Statistics Kruskal-Wallace One-way ANOVA versus naïve medium, n = 4 biological replicates

**Figure S4**

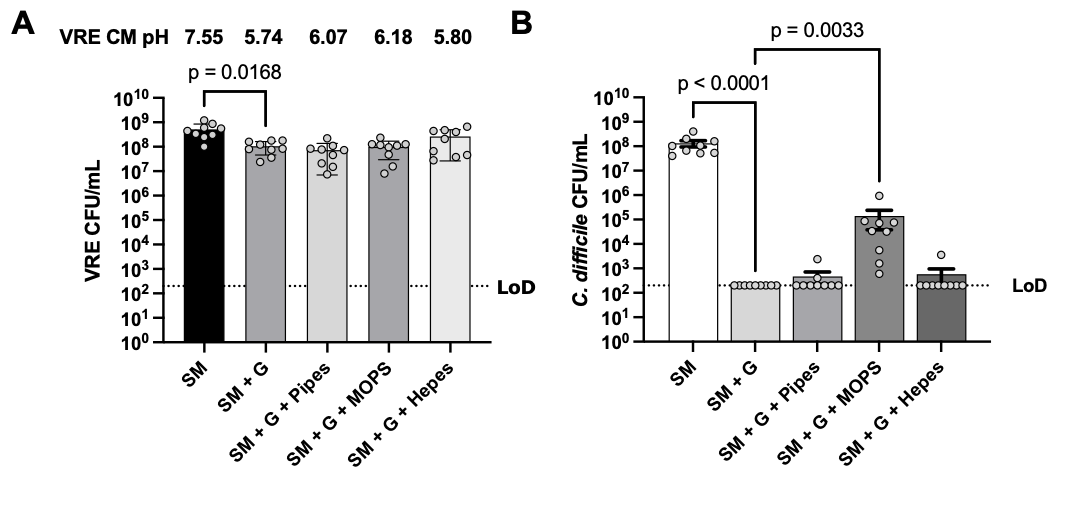

**Supplementary Figure 4:** **Buffering partially restores *C. difficile* growth in high glucose conditioned media.** A) VRE was grown for 24 hours in SM liquid medium with and without 100mM buffers as labeled on the x-axis (G = 0.6% glucose), and plated for VRE. Medium was filter sterilized, pH was measured and used as growth medium for *C. difficile* in Figure S1B. Data are combined from 3 independent experiments, n = 9 biological replicates, Kruskal-Wallace one-way ANOVA with Dunn’s correction versus SM + G B) *C. difficile* CFU after 48 hours growth in VRE-conditioned medium, Kruskal-Wallace one-way ANOVA with Dunn’s correction versus SM + G.

**Table of Strains**

**Table S1: Table of Strains**

| **Species** | **Strain** | **Source** |
| --- | --- | --- |
| *Clostridioides difficile* | VPI 10463 | ATCC |
| *Enterococcus faecium* (VRE) | 700223 | ATCC |
| *Enterococcus hirae* | 9790 | ATCC |
| *Enterococcus faecium* | 19434 | ATCC |
| *Enterococcus faecalis* | 19433 | ATCC |
| *Clostridioides difficile* | R20291 | DSMZ |
| *Paraclostridium bifermentans* | 638 | ATCC |
| *Clostridium scindens* | 35704 | ATCC |
| *Clostridium citronae* | BAA-1317 | ATCC |
| *Clostridium innocuum* | 14501 | ATCC |
